# The Expression of Vesicular Glutamate Transporter 1 (VGLUT1) in the Rat Larynx and Implications for Laryngeal Proprioception

**DOI:** 10.1101/2023.03.02.530889

**Authors:** Victoria X. Yu, Ignacio Hernández-Morato, Susan Brenner-Morton, Michael J. Pitman

## Abstract

Proprioception plays a crucial role in laryngeal function. Further, dysfunctional proprioception likely contributes to disorders such as laryngeal dystonia, dysphagia and vocal fold paresis. Despite this, the physiology of laryngeal proprioception is not well-understood. Controversy remains over whether canonical proprioceptive organs, like muscle spindles (MS) even exist in the intrinsic laryngeal muscles (ILM). Vesicular Glutamate Transporter 1 (VGLUT1) expression has been described in the sensory afferents of MS. This study’s primary aim is to determine whether the ILM contain MS using VGLUT1. This is a novel approach, as prior studies have relied on morphology and myosin composition to study this question. Secondarily, we describe the pattern of VGLUT1 expression in the rat larynx, Larynges of 62 Sprague-Dawley rats distributed across 5 age groups (P3, P8, P11, P14-15, and adult), were sectioned and immunostained for VGLUT1 and beta-tubulin III. Other markers (S46, GNAT3, PLCβ2, S100b, CGRP) were used to further characterize identified afferent innervation. Of 62 rats, MS were identified in the lateral thyroarytenoid muscles of just three P8 rats, and no golgi tendon organs (GTO) were seen. VGLUT1-positive intramuscular receptor-like entities were observed ILM, and VGLUT1-positive nerve endings were observed in the laryngeal mucosa, concentrated around the arytenoid cartilage. Employing VGLUT1 immunostaining, this study shows that rat intrinsic laryngeal muscles rarely express MS and do not express GTO. This leaves open the possibility that the larynx exhibits a unique proprioceptive apparatus. VGLUT1-positive intramuscular and mucosal structures provide candidates for an alternative system. Further defining the role of these sensory organs will increase our understanding of vocal fold function and ultimately lead to better treatment of vocal fold disorders.

**KEY POINTS:**

1. Dysfunctional laryngeal proprioception likely contributes to disorders such as laryngeal dystonia, dysphagia, and vocal fold paresis. Unlike proprioception of skeletal muscles, proprioception of the intrinsic laryngeal muscles is poorly understood.
2. In the present study we demonstrate that canonical proprioceptive organs (muscles spindles and Golgi tendon organs) are rarely expressed in the rat larynx, by studying the expression pattern of VGLUT1.
3. We also demonstrate the presence of other sensory innervation and structures which may contribute to an alternative proprioceptive circuitry, which requires further study.

## INTRODUCTION

The larynx is an organ of the upper respiratory tract that participates primarily in respiration, deglutition, and phonation (Sasaki, 2006; McHanwell, 2008). These functions, phonation in particular, involve near-instantaneous microadjustments of vocal fold position by the intrinsic laryngeal muscles (ILM). The coordination of the muscles to achieve such meticulous control of the glottic aperture requires fine-tuned proprioception (Hisa et al., 1985; Sasaki, 2006).

The physiology of proprioception in skeletal muscles is well-understood. Proprioception is carried out by two types of canonical proprioceptive end organs: muscle spindles (MS) and Golgi tendon organs (GTO). MS are encapsulated bundles of modified muscle fibers called intrafusal fibers, which are embedded within a muscle. MS are innervated by myelinated afferent Group Ia and Group II nerve fibers. Sensory information is projected through these fibers to the central nervous system, and synapses onto alpha and gamma motor effectors. (Golgi, 1880; Pedrosa-Domellöf et al., 1995; Jacobson and Marcus, 2008; Brushart, 2011).

By contrast, the physiology of proprioception of head and neck muscles, which are innervated by cranial nerves, is far less well-understood. MS have been found in masseter muscle, yet identification of canonical proprioceptive organs in other head and neck muscles remains elusive (Dessem et al., 1997; Rosales and Dressler, 2010). The presence of MS in the ILM in particular is controversial. While morphological studies have reported the presence of MS in the ILM, studies of ILM myosin heavy chain composition refute this in all but the interarytenoid muscle (Keene, 1961; Baken, 1971; Larson et al., 1974; Desaki et al., 1997; Sanders et al., 1998; Brandon et al., 2003a; b; Tellis et al., 2004). This raises the question of whether or not canonical proprioceptive organs exist in the vocal folds or whether perhaps the larynx has an alternative proprioceptive apparatus.

The purpose of this study is to examine this question using a novel approach by investigating Vesicular Glutamate Transporter 1 (VGLUT1)-positive sensory innervation of the ILM. VGLUT1 belongs to a family of transporters (VGLUT1, 2, and 3) that facilitate packaging of neurotransmitter glutamate into presynaptic vesicles. The three VGLUT isoforms are differentially expressed in the central (CNS) and peripheral (PNS) nervous systems. VGLUT1 in particular has been implicated in mechanosensation. Centrally, VGLUT1 is expressed in lamella of the spinal cord associated with mechanosensation. Peripherally, VGLUT1 is expressed in dorsal root ganglia (DRG) and mesencephalic trigeminal nucleus neurons associated with mechanosensation, in projections terminating on motoneurons in the trigeminal motor nucleus, and in nerve endings terminating on MS (Varoqui et al., 2002; Oliveira et al., 2003; Todd et al., 2003; Wu et al., 2004; Pang et al., 2006; Shneider et al., 2009; Lund et al., 2010; Alvarez et al., 2011; Bullinger et al., 2011; Zhang et al., 2018).

This is a novel approach, because prior studies have relied on MS morphology and intrafusal fiber myosin composition to identify proprioceptive organs. Using VGLUT1 and examining the question of laryngeal proprioception from an innervation standpoint broadens the scope of investigation to allow for the possibility of an alternative end organ. Our first aim is to settle the controversy over the presence of MS in the ILM, as VGLUT1 has a characteristic staining pattern within MS (Wu et al., 2004; Pang et al., 2006; Shneider et al., 2009; Lund et al., 2010). The second aim is to evaluate other sensory structures in the vocal folds.

## MATERIALS AND METHODS

### Experimental Animals

Sixty-two Sprague Dawley rats were used in this study. Animals were distributed across 5 age groups: postbirth days 3 (P3), 8 (P8), 11 (P8), and 14/15 (P14/15), and 60-day-old young adults (Table 1). All procedures were performed in accordance with the Public Health Service Policy on Humane Care and Use of Laboratory Animals and the Animal Welfare Act (7 U.S.C. et seq.). Columbia University’s Institutional Animal Care and Use Committee approved all protocols and animal handling.

**Table 1.**
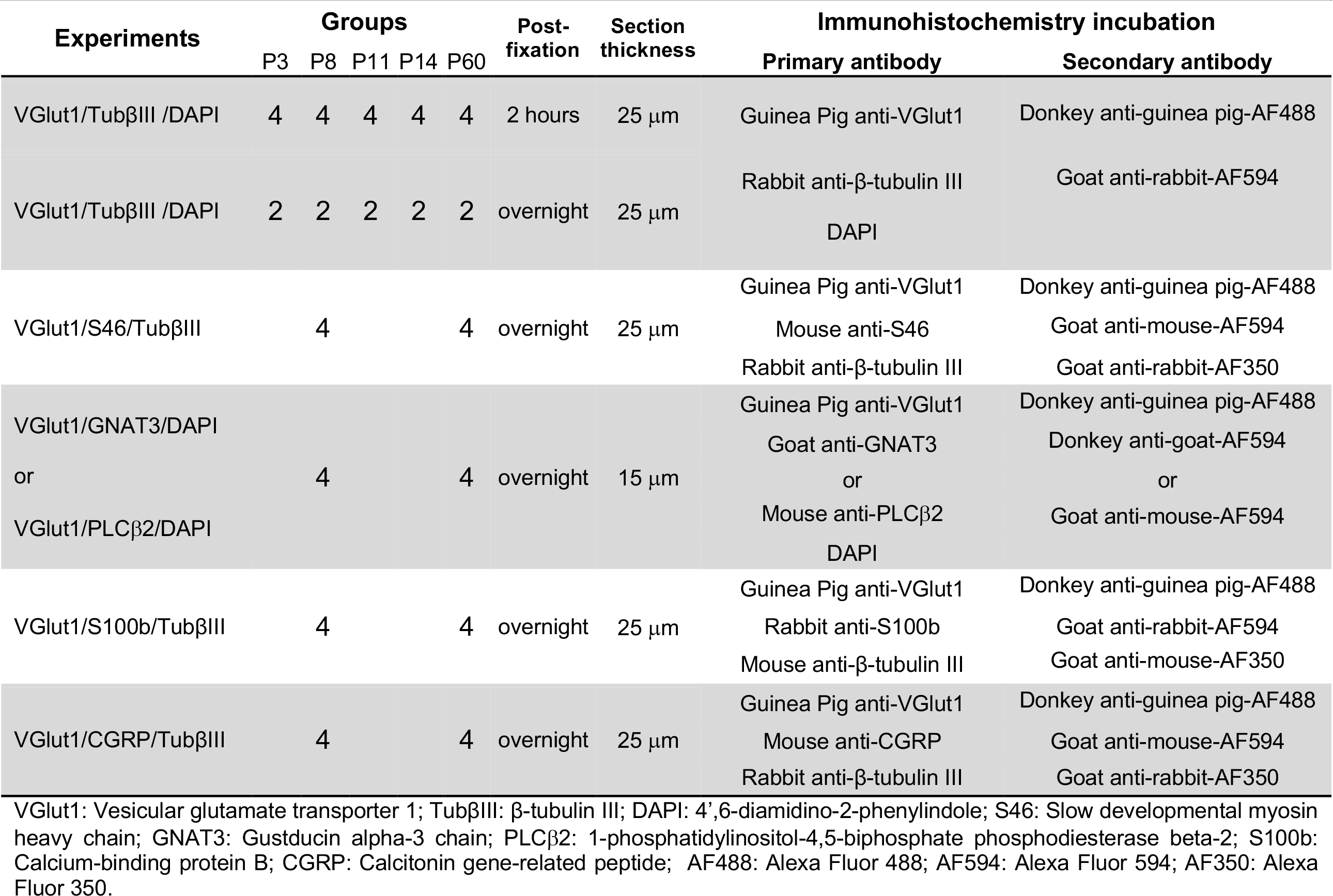
Number of animals used in this study.

#### Tissue Preparation

Animals were euthanized with a lethal dose of ketamine (100 mg/kg) and xylazine (10 mg/kg). They were then transcardially perfused with sterile saline solution, followed by 4% paraformaldehyde in phosphate buffer saline (PBS). Because some of the studied structures were difficult to identify, good perfusion was essential for removing blood cells, which could be misinterpreted as positive findings. Larynges were then dissected out. Two post-fixation protocols were tested as outlined in Table 1: larynges post-fixed for two hours and larynges postfixed overnight in 4% paraformaldehyde at 4ºC. Next, all the larynges were immersed in 15% sucrose in PBS, followed by 30% sucrose in PBS for cyroprotection. They were then embedded in OCT Compound (Sakura, Torrance, CA) and stored at -80ºC. Hindlimb muscles (gastrocnemius, extensor digitorum longus, soleus) were also processed for use as control samples.

### Experimental groups

First, a set of 30 animals was used to observe the pattern of VGLUT1 staining in the rat larynx. Subsequently, several experiments were performed to study colocalization of VGLUT1 with various other markers: 1) a set of eight animals to study colocalization of VGLUT1 and marker of intrafusal fibers S46, 2) a set of eight animals to study colocalization of VGLUT1 and gustatory markers GNAT3 and PLC beta 2, 3) a set of eight animals to study VLUGT1 and myelination marker S100b, and 4) a set of eight animals to study VGLUT1 and nociceptor marker CGRP.

### Immunohistochemistry of rat larynges

Larynges were sectioned coronally using a cryostat in several thicknesses as indicated in Table 1. Sections were sequentially captured onto gelatin-coated slides.

The primary and secondary antibodies used were summarized in Table 2 and 3. Guinea pig anti-VGLUT1 antibody was kindly provided by Susan Morton. Prior to staining, sections were left at room temperature for one hour and then washed in PBS twice for five minutes. Experimental groups staining for S46, CGRP, GNAT3, and PLCβ2 required preincubation in goat (S46, CGRP, PLCβ2) and donkey (GNAT3) serum for primary antibody incubation, which was performed with 5% serum in PBS with 0.3% Triton (PBST) for 30 minutes. Primary antibodies incubation was performed in PBST for 48 hours at 4°C. For experiments requiring preincubation for secondary antibody incubation, 2% serum was added. Then, slides were washed once and secondary antibody staining was done for two hours at room temperature. Hindlimb muscles and extrinsic laryngeal muscles attached to the larynx served as control samples.

**Table 2.**
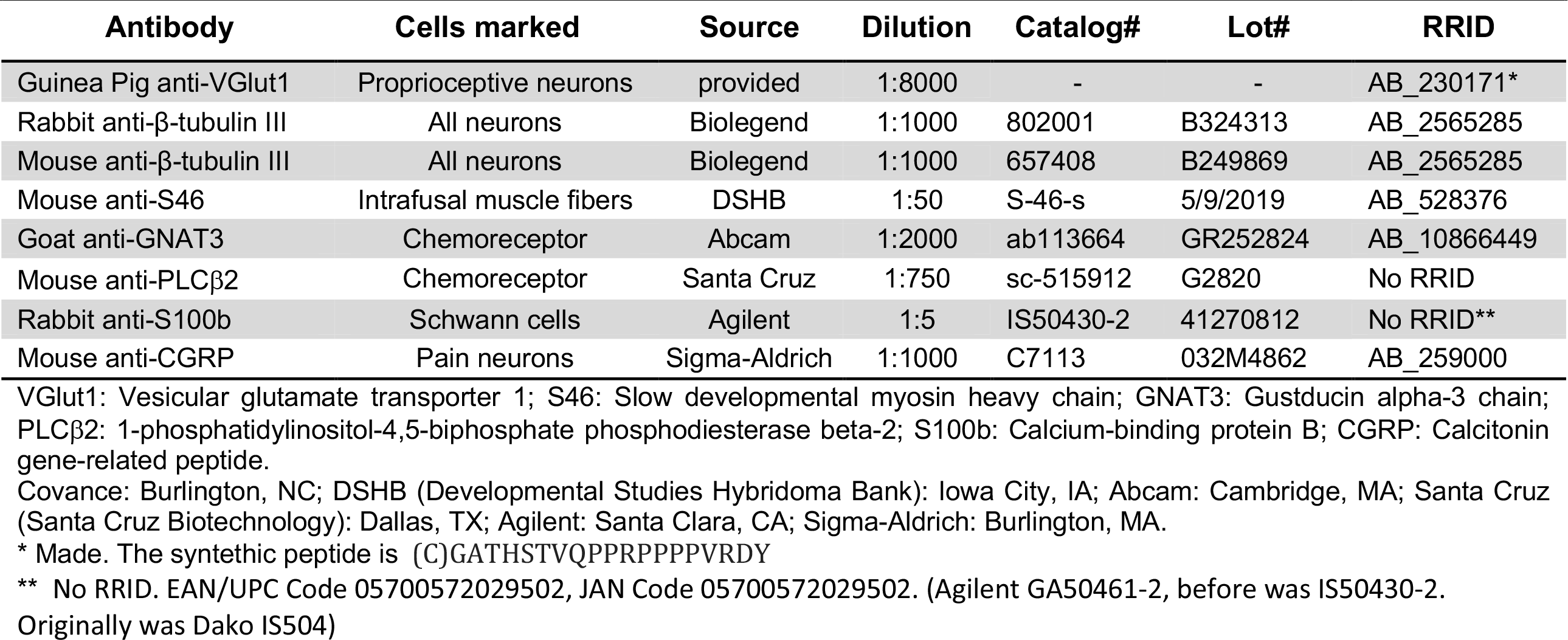
Primary antibodies.

**Table 3.**
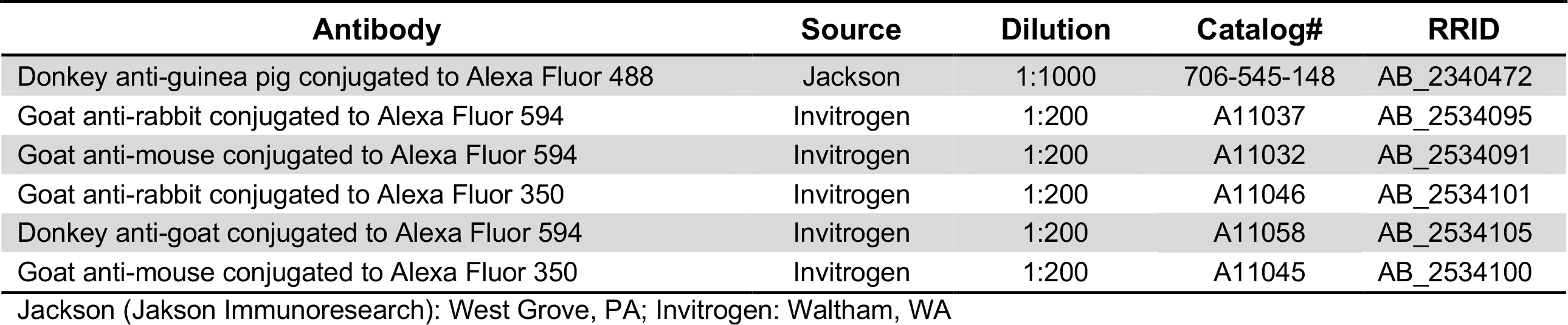
Secondary antibodies.

Coverslips were mounted with Fluoroshield Mounting Medium with 4’,6-diamidino-2-phenylindole (DAPI) (Abcam) and stored at 4ºC. When larynges were stained with S46, S100b, and CGRP mounting medium without DAPI was used (Sigma). All samples were analyzed using a Zeiss Axioskop epi-fluorescence microscope (Zeiss, Oberkochen, Germany). Experiments were analyzed at the microscope by two researchers (VY and IH) for MS based on previously described staining patterns in the literature, which were reproduced in our control tissues. IH also evaluated laryngeal and hindlimb tissue for GTO based on previously described staining patterns (Woo et al., 2015). In contrast, the flower-spray like endings, intramuscular receptors, and mucosal receptors were novel findings that required consensus building through several rounds of reassessment of tissue samples, not allowing for two-observer independent review. The following criteria for identifying each of these structures were ultimately decided on and will be used in future studies to allow independent review: 1) Intramuscular receptors: VGLUT1-positive bulbous entities, typically located in muscle bellies and individually innervated by VGLUT1-positive medium to large nerve fibers; 2) flower spray-like endings: VGLUT1-positive nerve fibers that branch and form varicose terminals which splay over the surface of a muscle fiber; 3) mucosal receptors: VGLUT1-positive conglomerations below the epithelial layer at the laryngeal mucosa and innervated by VGLUT1-positive nerve fibers. Consensus findings for each animal are listed in Supplemental Tables 1 and 2.

The same slides were used to assess all 4 types of structures. We were unable to mask for structure and muscle when examining slides, as the shape of these entities are recognizable to the educated observer.

## RESULTS

### Muscle spindles

Two tissue controls were used: hindlimb muscle and extralaryngeal muscles, both of which are known to contain MS. MS were visualized in both types of control tissue using VGLUT1 and beta tubulin III at every time point (Figure 1 and 2). Beta tubulin III is a microtubule protein expressed in neurons (Tischfield et al., 2010).

**Figure 1.**
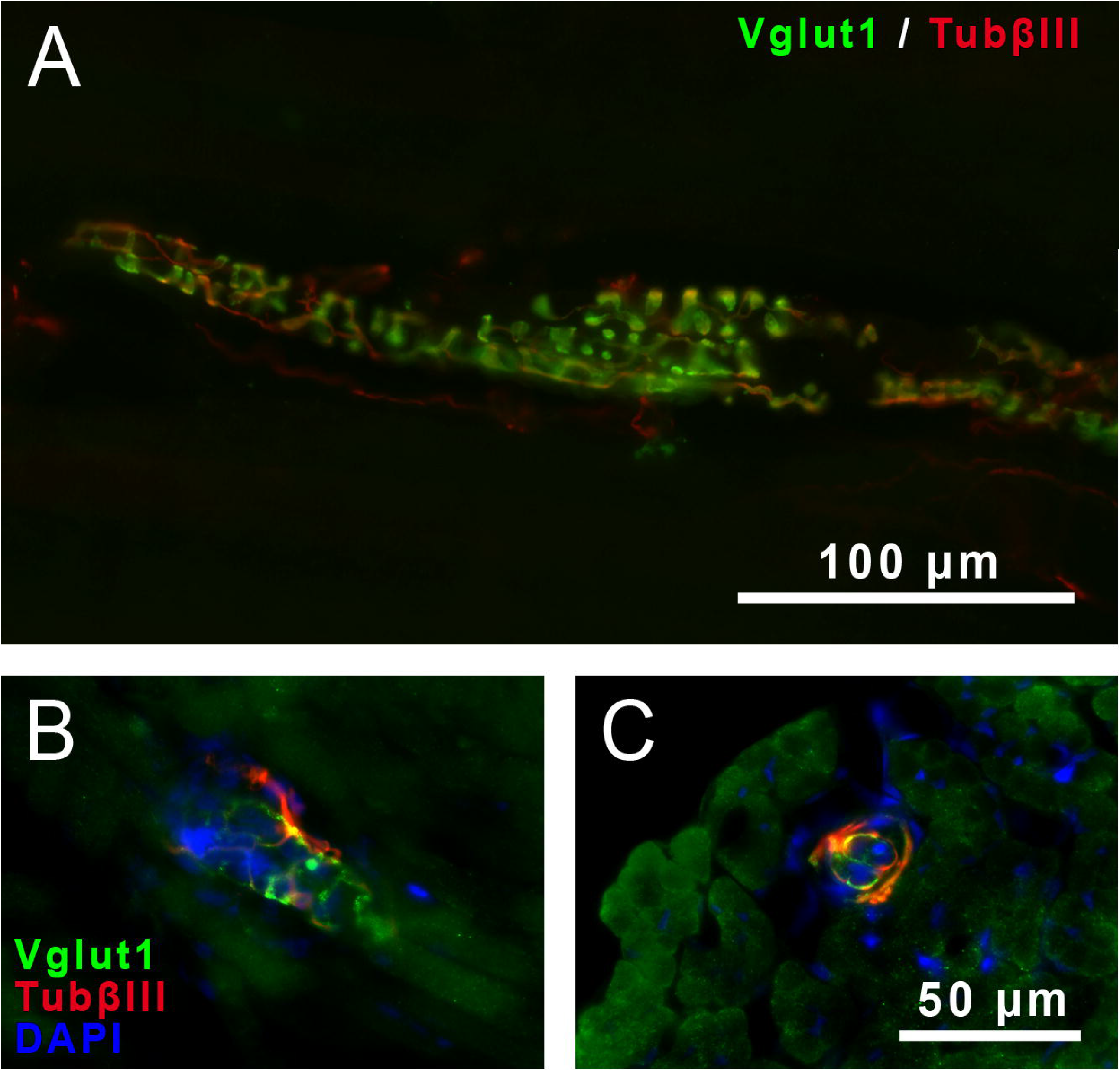
Respresentative muscle spindle identified by immunostaining for VGLUT1 and Tubulin-βIII in the gastrocnemius muscles (control tissue) in a P14 rat (**A**). Muscles spindles also identified in the extralaryngeal muscles of a P3 rat (**B, C**).

**Figure 2.**
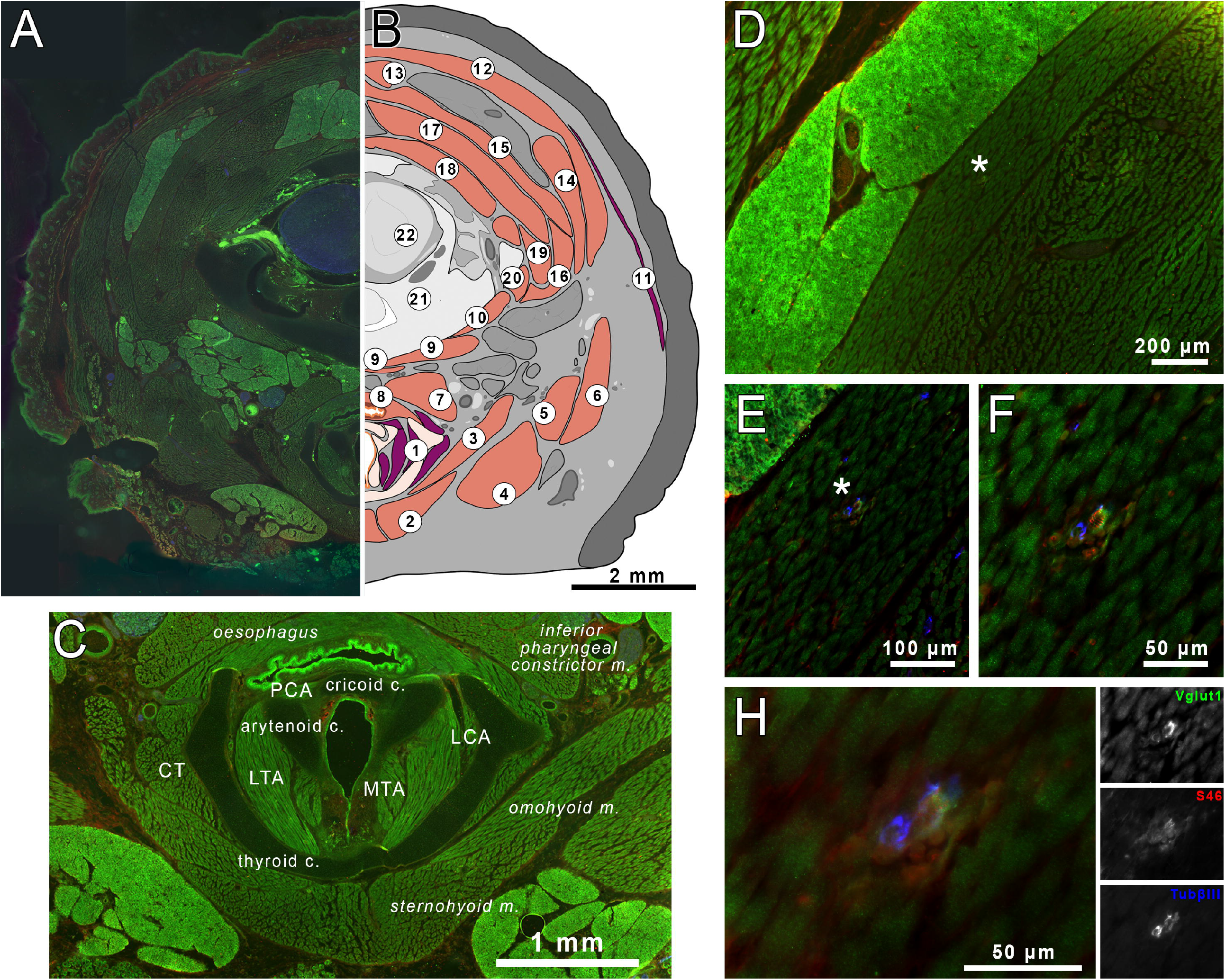
Muscle spindles identifies in neck muscles at the level of the larynx. Composition of several images taken a 5X of the neck at P0 rat (A). A diagram showing muscles of the neck; muscles in which muscle spindles were found are in pink; muscles in which muscle spindles were absent are in purple (B). A composite of 5X images of the larynx to include intrinsic and extrinsic laryngeal muscles (C). Muscle spindles stained for VGLUT1 (green), S46 (red) and Tubulin-βIII (blue) within neck muscles were observed using 4X magnification (D), 10X (E), 20X magnification (F), and 40X (G). Legend for diagram in panel B: 1 – larynx, 2 – sternohyoid muscle, 3 – omohyoid muscle, 4 – posterior portion of the digastric muscle, 5 – sternomastoideus muscle, 6 – cleidomastoideus muscle, 7 – inferior pharyngeal constrictor muscle, 8 – oesophagus, 9 – longus colli, 10 – longus capitis, 11 – platysma, 12 – cervicoauricularis, 13 – trapezius, 14 – rhomboideus capitis, 15 – splenius, 16 – rectus capitis lateralis, 17 – rectus capitis dorsalis, 18 – semispinalis cervicis, 19 – longissimus capitis, 20 – longissimus atlantis, 21 – cervical vertebrae, 22 – spinal cord, LTA – lateral thyroarytenoid muscle, MTA – medial thyroarytenoid muscle, LCA – lateral cricoarytenoid muscle, CT – cricothyroid muscle, PCA – posterior cricoarytenoid muscle.

In contrast, rare MS were observed in the ILM (Figure 3, Figure 4F and G). Two experimental conditions were used for this set of experiments, varying the postfixation time [2 hours postfix (n=20) and overnight postfix (n=42)] to rule out postfixation time as a confounder. Between both groups, MS were observed in the lateral thyroarytenoid muscles of three of 62 rats (Figures 3, 4F and G). All three rats were P8. In two rats, MS were observed unilaterally, and in one rat, bilaterally. VY and IH performed independent review of samples for MS, with concordant findings.

**Figure 3.**
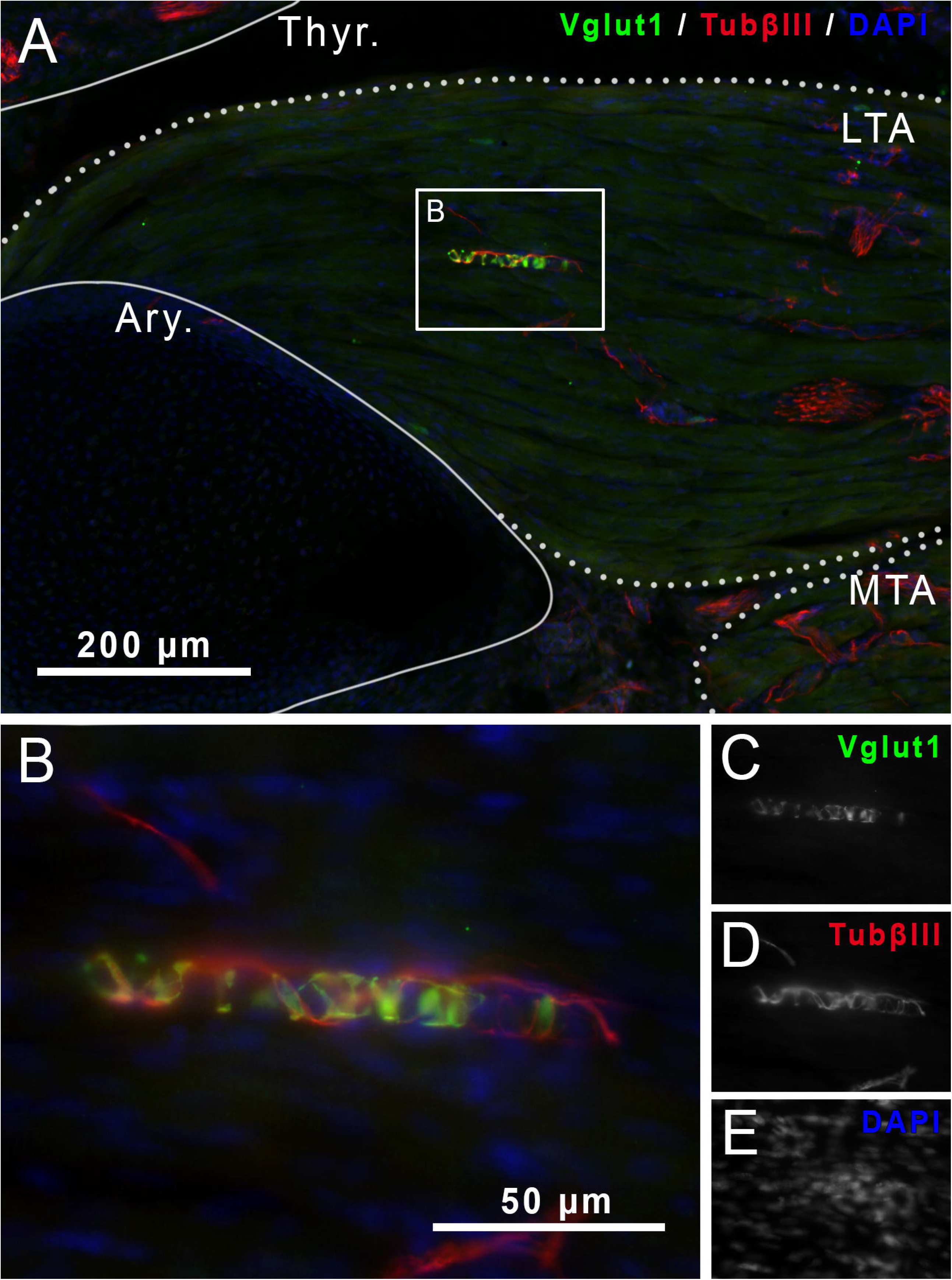
Muscle spindle in the lateral thyroarytenoid muscle of a P8 rat (**A-B**) with characteristic VGLUT1 staining (green, **C**) that colocalizes with Tubulin-βIII (red, **D**). DAPI in blue (**E**). Thyr. Thyroid cartilage, Ary. Arytenoid cartilage, LTA. Lateral thyroarytenoid muscle, MTA. Medial thyroarytenoid muscle.

**Figure 4.**
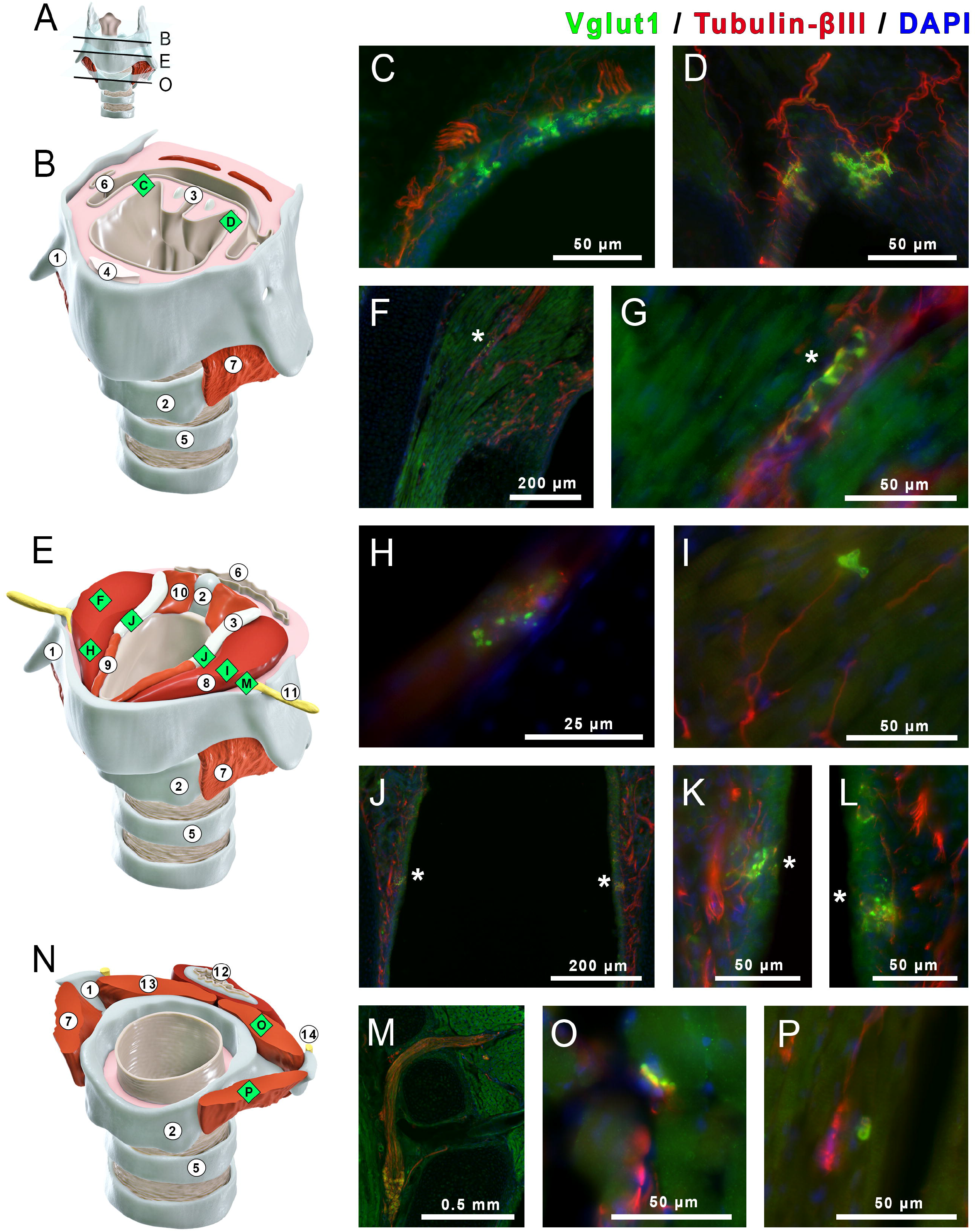
Immunohistological findings at various levels of the rat larynx. In (**A**), diagrammatic representation of rat larynx labels at three levels: supraglottis (**B**), glottis at the level of the vocal folds (**E**), and subglottis (**N**). In the supraglottis (**B)**, the only findings were mucosal formations in the aryepiglottic fold (**C, D**). In the glottis (**E**), the findings included muscles spindles in the lateral thyroarytenoid in 3 P8 rats (**F, G**), flower-spray endings (**H**), intramuscular receptor (**I**), mucosal receptors (**J-L**), internal branch of the SLN entry and parasympathetic intralaryngeal ganglia (**M**). In the subglottis (**N**), the findings were intramuscular receptors (**O**, in the posterior cricoarytenoid and **P**, in the cricothyroid). Structures of the larynx are numbered in the diagram: 1. Thyroid cartilage, 2. Cricoid cartilage, 3. Arytenoid cartilage, 4. Epiglottis, 5. Tracheal ring, 6. Pharynx, 7. Cricothyroid muscle, 8. Lateral thyroarytenoid muscle, 9. Medial thyroarytenoid muscle, 10. Superior cricoarytenoid muscle, 11. Internal branch of the superior laryngeal nerve, 12. Esophagus, 13. Posterior cricoarytenoid muscle, 14. Recurrent laryngeal nerve.

Additionally, of the 62 rats, we stained for S46 in four P8 and four adult rats. S46 is a slow-tonic myosin heavy-chain present in intrafusal bag fibers (Schiaffino and Reggiani, 2011). Hindlimb and extralaryngeal muscles were used as controls; MS in these tissues demonstrated colocalization of S46 and VGLUT1 (Figure 5C). No S46 staining was observed in any ILM of the eight animals used for the S46 set of experiments.

**Figure 5.**
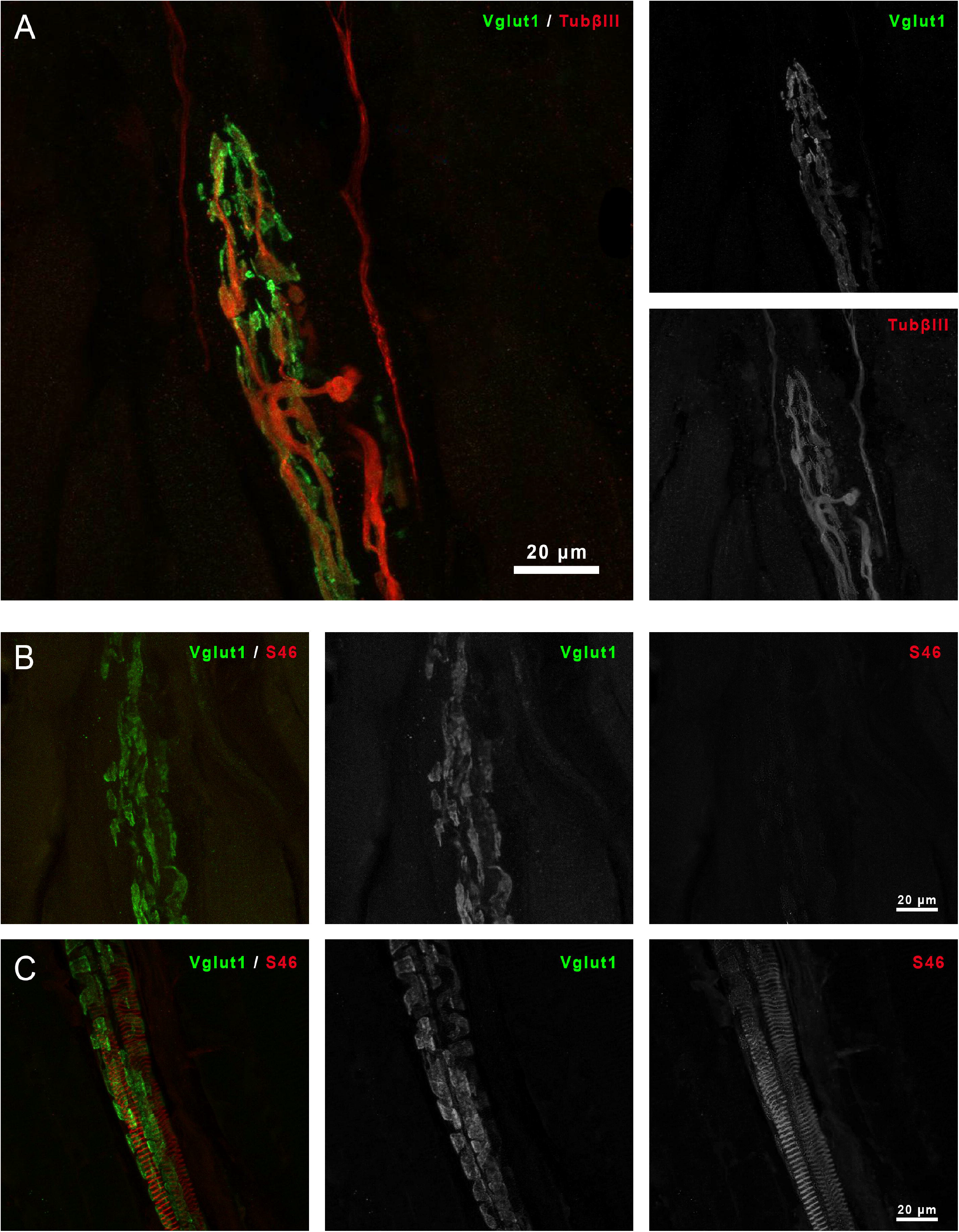
Golgi tendon organs in adult rat hindlimb stains positive for VGLUT1 and Tubulin-βIII (**A**). Golgi tendon organs are positive for VGLUT1 but negative for intrafusal fiber marker S46 (**B**). In contrast, muscles spindles stain positive for VLUGT1 and S46 (**C**).

### Golgi tendon organs

GTOs were noted in control hindlimb muscle but not in the ILM. Note that GTOs stain positive for VGLUT1 and beta tubulin III but, unlike MS, are negative for S46 (Figure 5A and 5B).

### Intramuscular receptors

After noting that MS were largely absent in the ILM, close examination of coronal sections of the larynx was performed to characterize other sensory innervation and structures. In a subset of 54 study animals, VGLUT1 and beta tubulin III co-staining revealed intramuscular receptor-like entities in the ILM of 30 animals across all age groups (Figure 6). They were comprised of medium-sized nerve endings terminating in bulbous formations on muscle fibers (Figure 6). Where present, one to five receptors were identified in each muscle. The location of these VGLUT1 positive bulbous formations within the muscle varied among the ILM. For instance, in the posterior cricoarytenoid muscle these entities were observed at the periphery, whereas, in the lateral thyroarytenoid muscle, they were identified deep in the muscle (Figure 4O and 4P). These entities were found to stain with S100b but not with CGRP (Figure 8). S100b is a calcium-binding protein expressed in myelin-producing Schwann cells (Mata et al., 1990). CGRP is a neuropeptide that is a key neurotransmitter in nociception (Yarwood et al., 2017).

**Figure 6.**
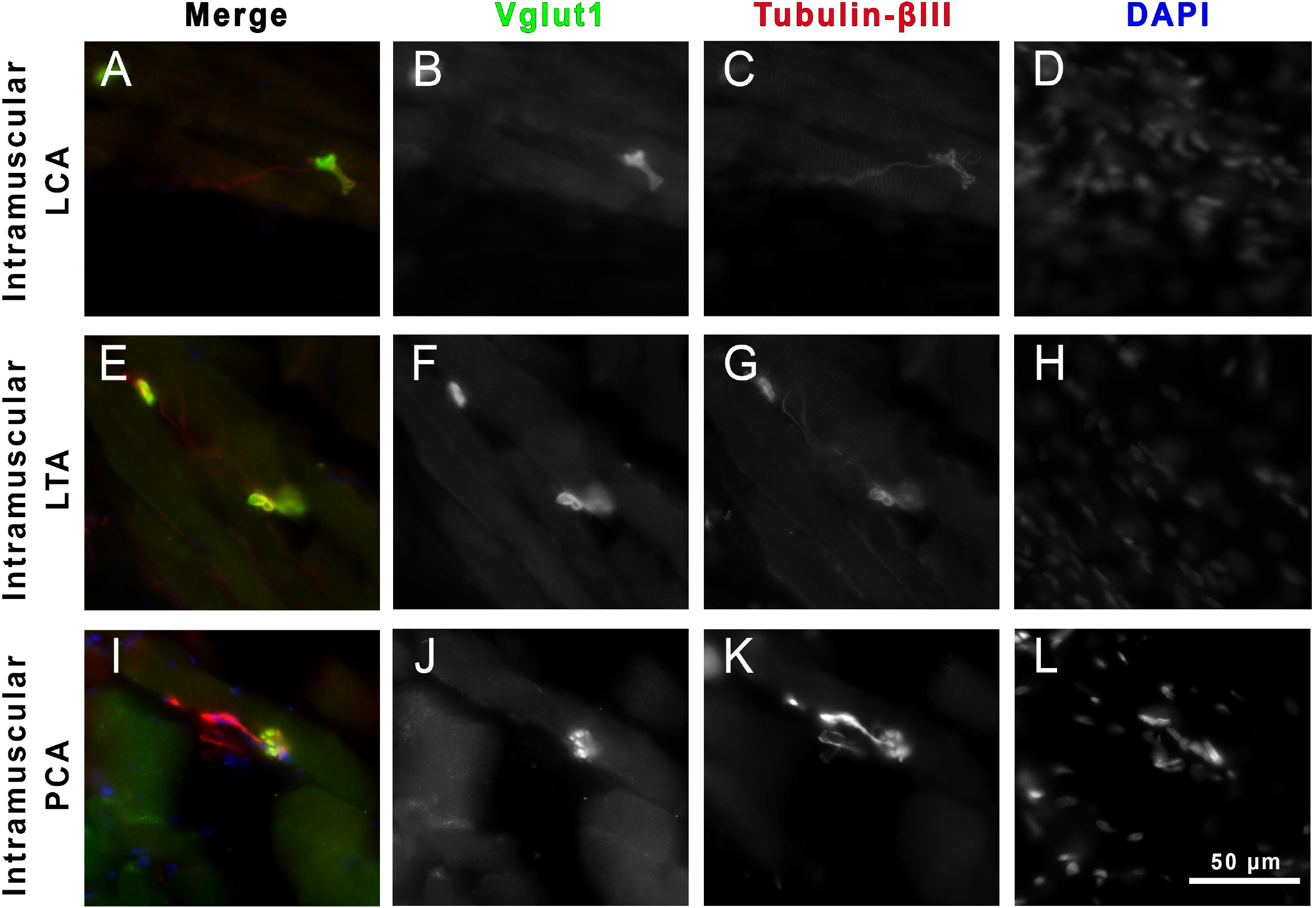
VGLUT1-positive intramuscular receptor-like entities in the lateral cricoarytenoid (LCA) (**A-D**), the lateral thyroarytenoid (LTA) (**E-H**), and the posterior cricoarytenoid (PCA) (**I-L**).

### Flower spray-like endings

In a subset of 54 study animals, VGLUT1-positive flower spray-like nerve endings were observed to terminate on extrafusal fibers in the lateral thyroarytenoid muscles of 5 rats, 3 P14 and 2 adults (Figure 7, Figure 4H). Like the intramuscular receptors, the flower spray-like endings were S100b-positive and CGRP-negative.

**Figure 7.**
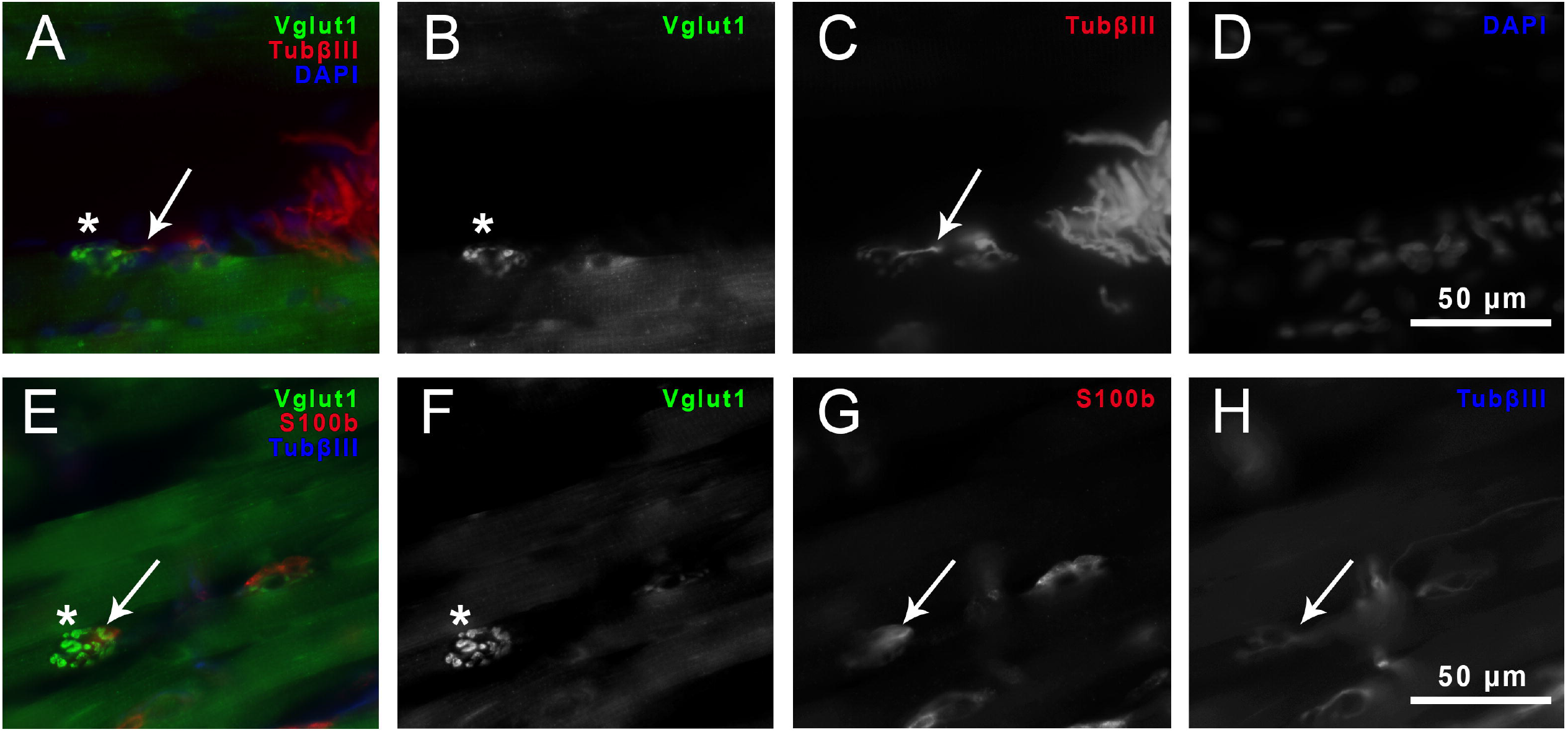
Flower-spray endings in the lateral thyroarytenoid muscle of a P14 rat (**A-D**) with characteristic VGLUT1 staining (green, **asterisks**) that colocalizes with Tubulin-βIII (red, **arrows**). Also, VGLUT1 (green, **asterisks**) colocalizes with S100b (red, **arrows**) at the flower-spray endings (**E-H**).

**Figure 8.**
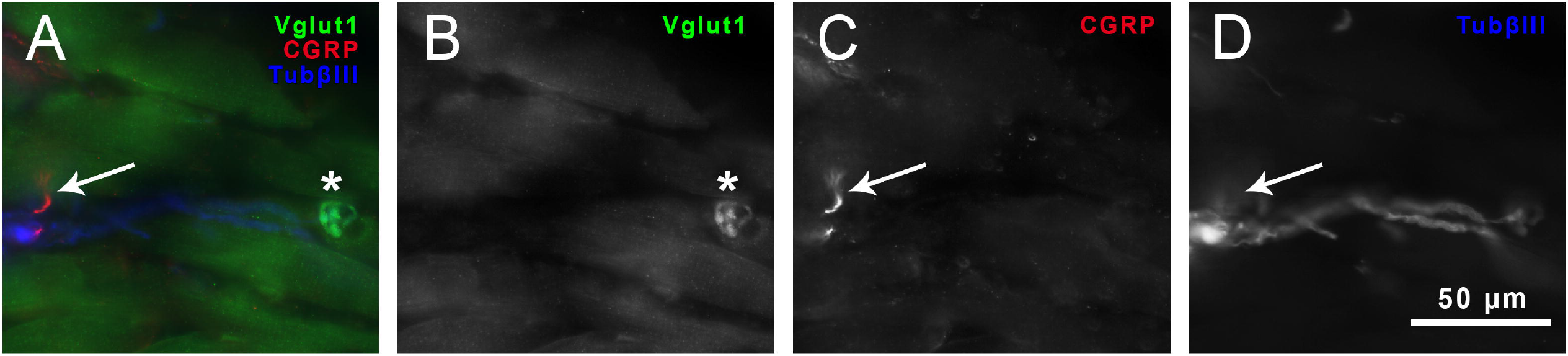
CGRP-positive nerve endings (red, **arrows**) do not colocalize to VGLUT1 formations (green, **asterisks**) at the intramuscular receptors (**A-D**).

### Mucosal receptors

Clear colocalization of VGLUT1 and beta tubulin III staining was observed in the laryngeal mucosa. Conglomerations of VGLUT1 expression were observed bilaterally in the mucosa of the aryepiglottic fold innervated by large-nerve-fibers at the supraglottic region (Figure 4C and 4D). VGLUT1 positive formations were also observed in the mucosa overlying the arytenoid cartilages at the glottis (Figure 4J, 4K, and 4L). These latter VGLUT1 foci were observed in several sections and appeared to run in two stripes longitudinally along the medial surface of the arytenoids. These mucosal formations were observed in all 62 rats, but VGLUT1 staining was strongest in the P3 group and weaker in the older groups.

GNAT3 and PLCβ2, markers of type II and III taste receptors, respectively, colocalized with VGLUT1 staining in some nerve fibers within the laryngeal mucosa (Figure 9E and 9F). However the arytenoid-associated VGLUT1-staining conglomerates were negative for both markers. Additionally, S100b staining was observed to colocalize with the arytenoid-associated VGLUT1-positive mucosal conglomerations, while they were CGRP negative (Figure 9A-9D). In contrast, clear CGRP-positive staining was identified in the laryngeal muscles, associated with small nerve fibers.

**Figure 9.**
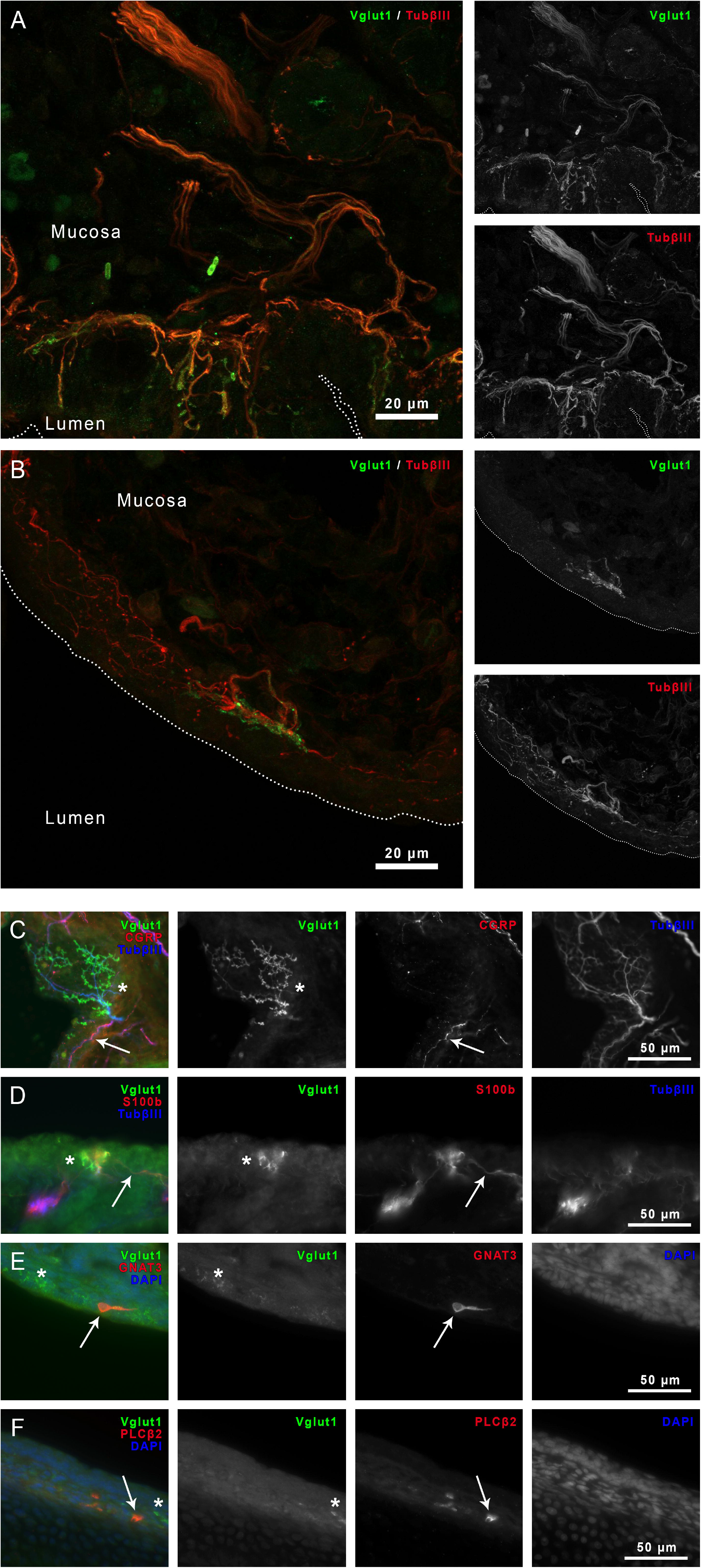
VGLUT1-positive nerve endings the laryngeal mucosa at the aryepiglottic fold (**A**) and along the medial surface of the arytenoids (**B**). CGRP-positive nerve endings (red, **arrows**) do not colocalize to VGLUT1 formations (green, **asterisks**) at the laryngeal mucosa (**C**). However, VGLUT1 (green, **asterisks**) colocalizes with S100b (red, **arrows**) at the laryngeal mucosa (**A-D**). VGLUT1-positive nerve endings at the laryngeal mucosa (green, **asterisks**) do not colocalize with taste receptors (red, **arrows**). GNAT3 (**E**), PLCB2 (**F**), taste receptor markers.

## DISCUSSION

The present study uses a novel approach of investigating proprioceptive end organs of the rat larynx by examining the expression pattern of VGLUT1 in the ILM. Previous studies have investigated the larynx for the identification of gross structures of MS or intrafusal fibers. These studies have reached conflicting conclusions regarding the presence of MS in the vocal folds, exemplified by investigations of the most studied ILM, thyroarytenoid muscle (TA). Studies from the 1950s to 1990s used traditional stains (H&E, silver, trichrome) to identify MS by morphology. Multiple groups described MS in the thyroarytenoid muscle (TA), with descriptions of MS concentrated in the superior medial quadrant of the TA in close proximity to the vocal ligament (Keene, 1961; Konig and Leden, 1961; Rossi and Cortesina, 1965; Katto et al., 1987; Sanders et al., 1998; Baken and Orlikoff, 2000). Others using these techniques reported no MS in the TA (Fernand and Young, 1951; Murray, 1957). Studies in the 2000s using IHC against MHC also refute the presence of MS in the TA (Brandon et al., 2003b).

Prior authors have provided possible explanations for these discrepancies, observing that ILM MS may be structurally different from stereotypical MS with features that include thinner capsules, unique chain fiber features, shorter intrafusal fibers, fewer fibers per MS, and/or branching complexes (Katto et al., 1987; DelGaudio et al., 1995; Desaki et al., 1997; 1998; Sanders et al., 1998; Tellis et al., 2004; Desaki and Nishida, 2006). Further and more generally, cranial muscles have been noted to have unique extrafusal fiber isoforms (DelGaudio et al., 1995).

Given the potential for ILM MS to exhibit non-canonical muscle and capsule structure and composition, we turn to innervation as a possible common thread and a different approach to determining whether MS exist in the ILM. We chose to study VGLUT1, a glutamate transporter, specifically. Centrally, VGLUT1 is expressed in excitatory glutamanergic neurons of several structures including the telencephalon, cerebellum, precerebellar nuclei, and medial habenular nucleus (Zhang et al., 2018). Importantly, VGLUT1 is expressed in axons terminating in regions of the spinal cord associated with low threshold cutaneous and proprioceptive afferents (deep dorsal horn, central canal, intermediate area, and ventral horn). These axons are also negative for markers of nociception, motor, and interneurons, and when dorsal root rhizotomy is performed, VGLUT1 staining density decreases (Todd et al., 2003; Wu et al., 2004; Brumovsky et al., 2007). This is in keeping with prior observations that central synapses of primary afferents, including group 1a afferents, are glutamanergic (De Biasi and Rustioni, 1988; Maxwell et al., 1990; Broman et al., 1993; Ornung et al., 1995; Banks et al., 2002; Brumovsky et al., 2007). Additionally, VGLUT1 is expressed in cell bodies of trigeminal mesencephalic nucleus neurons and in synapses on trigeminal motor nucleus motoneurons traced from the masseter muscle (Pang et al., 2006).

Peripherally, VGLUT1 is expressed in primarily medium and large, CGRP-negative dorsal root ganglia (DRG) neurons and fibers (Oliveira et al., 2003; Brumovsky et al., 2007). Additionally, VGLUT1 is expressed in MS in a characteristic pattern in limb and masseter muscles (Wu et al., 2004; Pang et al., 2006; Shneider et al., 2009; Lund et al., 2010). VGLUT1 expression has also been described in free nerve endings in the skin, follicular neural network B, and in Merkel cell neurite complexes of the palate and whisker pads (Hitchcock et al., 2004; Nunzi et al., 2004; Brumovsky et al., 2007). The role of VGLUT1 in the innervation of these end organs is not well understood, though theories include autogenic excitation of sensory endings and release of glutamate from Merkel cells for mechanosensory transduction (Wu et al., 2004; Higashikawa et al., 2019).

The main finding of this study is that based on staining with VGLUT1, MS were absent in the vocal folds of all but three of the 62 experimental animals. Co-staining with S46 confirmed this. These findings suggest that while MS are generally absent from rat vocal folds, they may occur, and their presence is rare and variable. This may explain the disparate findings in past research and suggest inter- and intra-species variability (Keene, 1961; Desaki et al., 1997; 1998; Brandon et al., 2003a; b; Tellis et al., 2004; Desaki and Nishida, 2006). Still, this finding of infrequent, seemingly sporadic MS is perplexing.

Outside of the masseter muscle, other head and neck muscles that are also derived from cranial mesoderm during development exhibit lack of MS (Dessem et al., 1997; Rosales and Dressler, 2010). An explanation could rest in the origin of the myoblasts for different muscles during development (Nathan et al., 2008; Shih et al., 2008). Specifically, the pharyngeal arches from which head and neck muscles are derived may encode non-canonical proprioceptive organs during cranial mesoderm myogenesis, including those for the vocal fold movement (Kelly, 2010; Michailovici et al., 2015; Ziermann et al., 2018).

The observation that most rats have neither MS nor GTOs raises the question of whether an alternative proprioceptive mechanism is present in the larynx. An example of such an alternative proprioceptive system is observed in the extraocular muscles of the cat, which contain specialized sensory nerve ending configurations, palisade endings, that mediate proprioception (Büttner-Ennever et al., 2002; 2006; Zimmermann et al., 2013). Another example are the facial mimetic muscles, where a myocutaneous mechanoreceptive mechanism has been proposed in which muscle fibrils insert directly into collagen networks of the dermis without a superstructure (May and Bramke, 2021). A third example lies the in the superior constrictor in which corpuscle-like structures that stain positive for putative mechanoreceptors have been identified as possible alternative proprioceptors (de Carlos et al., 2013; Cobo et al., 2017).

The current study revealed several VGLUT1-positive findings: 1) VGLUT1-positive, CGRP-negative, S100B-positive intramuscular receptor-like structures in the intrinsic laryngeal muscles; 2) VGLUT1-positive, CGRP-negative, S100B-positive flower-spray-like endings on extrafusal fibers of the intrinsic laryngeal muscles; and 3) VGLUT1-positive arytenoid-associated mucosal conglomerates, a subset of which were GNAT3/PLCβ2- and CGRP-negative and S100B-positive. Though more studies must be done to define the roles of these structures, they are potential contributors to an alternative somatosensory system.

Previous electrophysiological studies of laryngeal muscles have detected the presence of mechanoreceptors within the laryngeal muscles, but the receptors themselves have yet to be identified and characterized (Sampson and Eyzaguirre, 1964; Abo-el-Enein, 1966; Storey, 1968; Widdicombe, 2001). Flower-spray endings are a type of sensory nerve ending that innervate intrafusal fibers (Hines, 1927; Hinsey, 1934; Liu et al., 2009; Vaughan et al., 2015). The presence of flower-spray-like innervating extrafusal fibers could suggest a specialized role of select extrafusal fibers in proprioception as was previously observed and suggested by Brandon et al in 2003 (Brandon et al., 2003b). Finally, previous, well-accepted electrophysiological studies suggest an active role of the mucosa for the control of vocal fold movement such as in the laryngeal adductor response (Sampson and Eyzaguirre, 1964; Storey, 1968; Sant’Ambrogio et al., 1983; 1991; Andreatta et al., 2002). The exact mechanism of the mucosal receptors observed in this study, such as pressure or stretch, requires more investigation.

A final consideration is the age-related findings in our data. Specifically, 1) MS were observed in 3 P8 rats only, 2) flower spray endings were noted in 3 P14 and 2 adult rats, and 3) the mucosal findings were strongest in P3 and weaker in older rats. Rats produce ultrasonic vocalizations that vary based on activity and state. Studies have demonstrated that the frequency, complexity, and duration of these vocalizations change with age. Specifically, aged rats have lower, less complex vocalizations. Infant rats also produce a special distress cry that older rats do not (Peterson et al., 2013; Stark et al., 2020; Lenell et al., 2021). It is possible that rats require different degrees or modes of sensory input from the vocal folds as they age, though this requires more study. Pang et al note age-related trends in VGLUT1 expression in their study of rat masseter. Specifically, VGLUT1 immunoreactivity in trigmeminal mesenteric neurons progressively decreased from infancy to adulthood. Conversely, VGLUT1 immunoreactively increased at trigeminal motor nucleus terminals and in MS with age (Pang et al., 2006).

A key limitation of the study relates to the nature of being the first study to characterize the distribution of VGLUT1 in the larynx. Identification of the VGLUT1 structures (flower-spray-like, intramuscular receptors, and mucosal receptors) required consensus building, which precluded fully independent review by VY and IH. In future studies, the criteria developed during this study will be used for identifying these three novel structures, so an independent review is achievable. Additionally, we were unable to investigate directly whether the discussed entities were mechanosensory in nature, as a suitable reagent for mechanoreceptor marker Piezo 2 in the rat was not available. Future studies using available mouse models are planned. Finally, another limitation of the study is choice of microscopy; confocal would have allowed for better resolution and characterization of the described structures. Ultimately, these are novel findings, and much work is needed to confirm and clarify the role of these VGLUT1-positive sensory components.

## CONCLUSION

Staining with VGLUT1 shows that MS appear rarely and sporadically in the rat intrinsic laryngeal muscles. GTO are not present. Intramuscular receptor-like entities, flower-spray-like endings, and VGLUT1 innervation in the peri-arytenoid mucosa were identified and require future investigation to determine whether they contribute to a unique proprioceptive mechanism in the larynx.

## ACKNOWLEDGEMENTS

No funding sources. Many thanks are owed to Dr. Joriene C. De Nooij and the Jessell lab for providing expertise and reagents, without which this study would not have been possible.

## Notes

**DISCLOSURES:** The authors have not conflicts of interest to disclose.

### Competing Interest Statement

The authors have declared no competing interest.

